# New Atg9 phosphorylation sites regulate autophagic trafficking in glia

**DOI:** 10.1101/2024.07.03.601894

**Authors:** Linfang Wang, Shuanglong Yi, Shiping Zhang, Yu-ting Tsai, Honglei Wang, Margaret S. Ho

**Author notes:** Corresponding author: Margaret S. Ho, Ph.D. These authors contributed equally (Linfang Wang, Shuanglong Yi, Shiping Zhang).

## Abstract

We previously identified a role for dAuxilin (dAux), the fly homolog of Cyclin G-associated kinase (GAK), in glial autophagy contributing to Parkinson’s disease (PD). To further dissect the mechanism, we present evidence here that lack of glial dAux enhanced the phosphorylation of the autophagy-related protein Atg9 at two newly identified threonine residues, T62 and T69. Enhanced Atg9 phosphorylation in the absence of glial dAux is potentially regulated through the master autophagy regulator Atg1, as the presence of which is required for Atg9 to interact with dAux in an otherwise separate condition. The enhanced Atg9 phosphorylation promotes autophagosome formation and Atg9 trafficking to the autophagosomes in glia. Whereas the expression of the non-phosphorylatable Atg9 variants suppresses the lack of dAux-induced increase in both autophagosome formation and Atg9 trafficking to autophagosome, the expression of the phophomimetic Atg9 variants restores the lack of Atg1-induced decrease in both events. Notably, Atg9 phosphorylation at T62 and T69 contributes to DA neurodegeneration and locomotor dysfunction implicated in PD. Thus, we have identified a dAux-Atg1-Atg9 axis relaying signals through the Atg9 phosphorylation at T62 and T69; these findings further elaborate the mechanism of dAux regulating glial autophagy and highlight the significance of protein degradation pathway in glia contributing to PD.

## Introduction

Glial cells are now recognized as important players in the neurodegenerative disease Parkinson’s disease (PD). As the second most common neurodegenerative disorder, PD patients exhibit classical motor symptoms accompanied by non-motor features, progressive degeneration of dopaminergic (DA) neurons, and the formation of Lewy bodies (LBs) ^1–3^. Glial cells contribute by releasing pro- or anti-inflammatory factors to exacerbate or ameliorate disease progression^4–8^, and signaling bidirectionally to regulate neuronal survival and function. Glial activation and intimate neuron-glia crosstalk are now indispensable features of PD onset and progression. Thus, unravelling the glial regulatory mechanism is crucial in advancing our understanding of PD.

Autophagy is a highly conserved catabolic process to engulf and degrade cytoplasmic materials and its dysregulation has been linked to many disorders including neurodegenerative diseases^9–13^. The activation of ATG1/Unc-51-like kinase 1 (ULK1) complex triggers the phagophore nucleation by phosphorylating components of the class III PI3K (PI3KC3) complex, leading to the local phosphatidylinositol-3-phosphate (PI3P) production and the recruitment of autophagy effectors DFCP1 and WIPI2. WIPI2 promotes LC3 lipidation via the ATG12-ATG5-ATG16L conjugation system for phagophore expansion, which then closes to form a double-membrane autophagosome. Subsequent fusion of autophagosome with the lysosome results in autolysosome, in which the contents are degraded and the salvaged nutrients are released back into the cytoplasm for recycling. Interestingly, the phagophore initiation and elongation are facilitated by ATG9, the sole transmembrane protein in the core autophagy machinery for providing lipids as autophagosome membrane sources^14,15^. It has been shown that ATG9 is transported to the pre-autophagosomal structure and autophagosomes to promote autophagy initiation upon autophagy induction^16–20^. ATG9 dysregulation has also been shown in α-syn overexpression or PD animal models, but how it contributes to PD remains largely elusive.

Here we present evidence that lack of dAuxilin (dAux), the *Drosophila* homolog of the PD risk factor Cyclin G-associated kinase (GAK, also known as DNAJC26), enhances Atg9 phosphorylation at two newly identified threonine resides T62 and T69 as revealed by the phophoproteomic analysis. The enhanced Atg9 phosphorylation promotes autophagosome formation and Atg9 trafficking to the autophagosomes. Whereas the expression of the non-phosphorylatable Atg9 variants suppresses the lack of dAux-induced increase in both autophagosome formation and Atg9 autophagic trafficking, the expression of the phophomimetic Atg9 variants restores the lack of Atg1-induced decrease in both events. Moreover, the Atg1 presence recruits Atg9 to interact with dAux. Considering that dAux interacts with Atg1 and Atg9 has been shown to be a Atg1 phosphorylation target, our results support the mechanism of a dAux-Atg1-Atg9 axis conveying signal via Atg9 phosphorylation at T62 and T69 for autophagy initiation in glia. Finally, enhanced Atg9 phosphorylation contributes to progressive DA neuron loss and locomotion deficits implicated in PD, demonstrating its pathological relevance.

## Results and Discussion

### Atg9 phosphorylation at T62 and T69 is upregulated in the absence of glial dAux

In our previous investigation, we uncovered the dAux-mediated autophagic mechanism in flies and its potential relevance to PD^24,25^. As we defined a link between dAux and the master regulator Atg1 in autophagy initiation, we sought to explore further how dAux confers its regulation on autophagy via Atg1. Revisiting the reported phosphoproteomic analysis detecting changes in phosphorylated peptides in 10-day-old adult fly heads of the control and *repo>daux*-RNAi flies^25^, we found that the phosphorylation at the two threonine residues: T62 (*daux*-RNAi/*LacZ* ratio 1.774) and T69 (*daux*-RNAi/*LacZ* ratio 1.774), of the autophagy protein Atg9 (the fly homolog of mammalian ATG9) was significantly upregulated upon glial dAux depletion (Figures 1A and 1B)^25^. Motif analysis predicted that T62 and T69 belong to sequence motifs prone to be phosphorylated, xxxxxx_T_ExExxx and xxxxxx_T_Pxxxxx, respectively (Figure 1C). T62, but not T69, is a highly conserved residue across species (Figure 1D), but neither of these two residues has been reported as targets for Atg9 phosphorylation. Given that lack of dAux causes enhanced phosphorylation of these two residues, Atg9 is unlikely to be a direct phosphorylation substrate of dAux.

**Figure 1.**
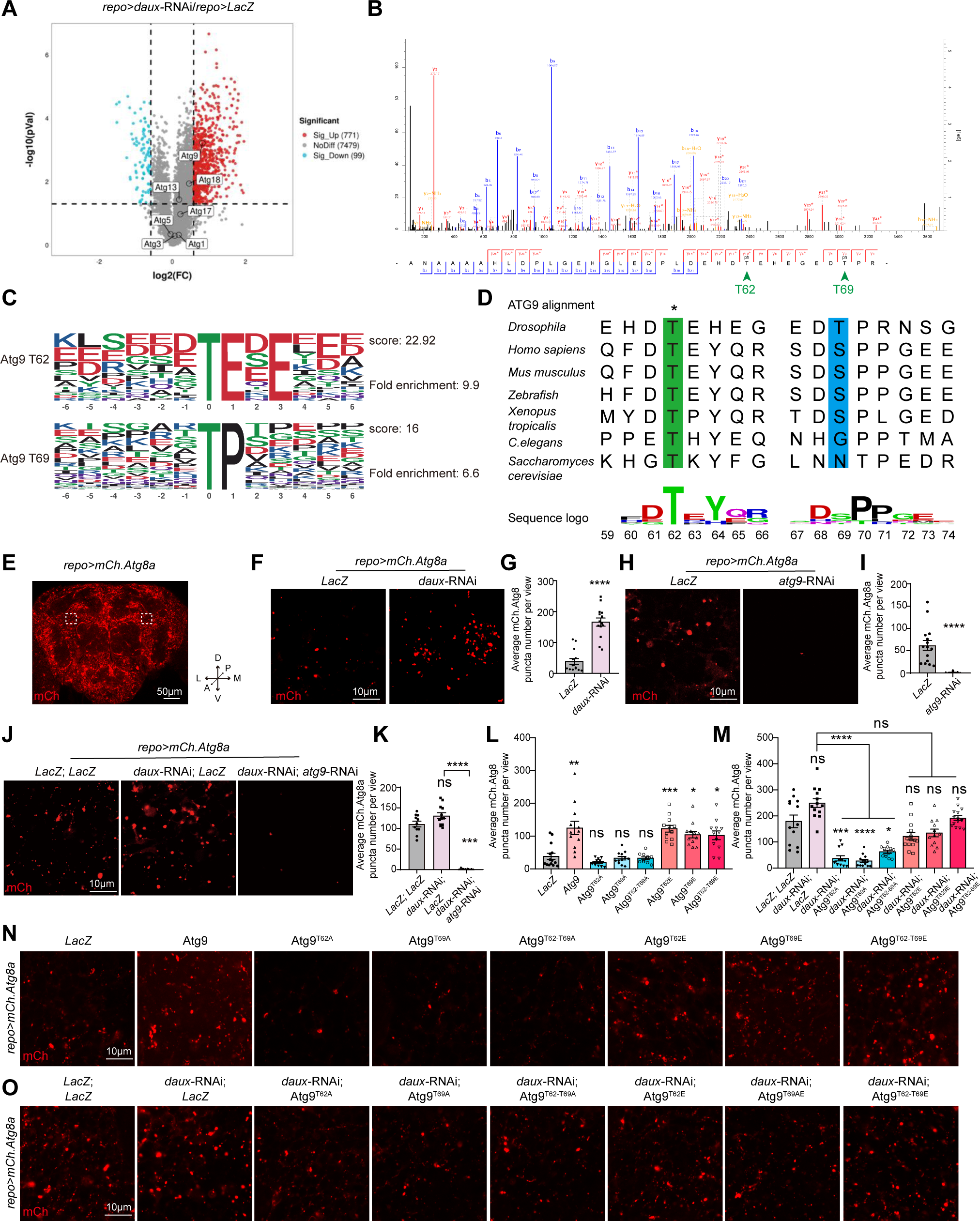
Lack of dAux-enhanced Atg9 phosphorylation at T62 and T69 promotes autophagosome formation in glia. (**A**) Advanced volcano plot drawn with phosphoproteomic data by the OmicStudio tools. The variance threshold is larger than 1.5 or less than 0.5, and p-value less than 0.05. (**B**) Phosphoproteomic analysis of the control and *repo>daux-*RNAi adult fly brains. Note that Atg9 phosphorylation at T62 and T69 are detected. (**C**) The sequences near T62 (xxxxxx_T_ExExxx) and T69 (xxxxxx_T_Pxxxxx) sites of Atg9 belong to phosphorylation motifs by the MoMo analysis tool using the motif-x algorithm. The motif consists of 6 amino acids upstream and downstream of the potential dAux modification site. Preferred amino acids are indicated with larger font, while less preferred amino acids are indicated with smaller font. (**D**) Alignment of Atg9 protein sequence across different species. Sequence logos are drawn with webLogo. Note that the T62 site is conserved across species. (**E**) An acquired microscopic image of an adult fly brain expressing *UAS-mCherry.Atg8a* under the control of the pan-glial driver *repo*-GAL4 (*repo>mCh.Atg8a*). The adult fly brain is positioned with the coordinates described (A: Anterior, P: Posterior, M: Medial, L: Lateral, D: Dorsal, V: Ventral), with the asymmetrical anterior/dorsal regions enclosed by white dotted squares selected for imaging in all Figures. (**F-K**) Representative images (F, H and J) and quantifications (G, I and K) of autophagosomes in adult fly glia. Note that the number of glial autophagosome increases and decreases respectively when expressing *daux*-RNAi (F and G) or *atg9*-RNAi (H and I). Co-expression of *atg9*-RNAi suppresses the *daux*-RNAi-induced increase in the autophagosome number (J and K). (**L-O**) Representative images (N and O) and quantifications (L and M) of autophagosomes in adult fly glia expressing different Atg9 variants with/without *daux*-RNAi. Note expressing Atg9 or either of the phosphomimetic Atg9 variants causes a significant increase in the glial autophagosome number, and expressing either of the three non-phosphorylatable Atg9 variants fails to cause significant difference (L and N). Expressing either of the three non-phosphorylatable Atg9 variants suppresses the increased autophagosome number in the absence of glial dAux, whereas expressing either of the three phosphomimetic Atg9 variants fails to reduce the increase significantly (M and O). Scale bars of different sizes are indicated on the images. Serial confocal Z-stack sections were taken at similar planes across all genotypes, with representative images shown as a single layer. Statistical graphs are shown with scatter dots indicating the number of brain samples analyzed. Data are shown as mean ± SEM. P-values of significance (indicated with asterisks, ns no significance, * p<0.05, ** p<0.01, *** p<0.001, and **** p<0.0001) are calculated by two-tailed unpaired t-test, Mann Whitney test, ordinary one-way ANOVA followed by Tukey’s multiple comparisons test, or Kruskal-Wallis tests followed by Dunn’s multiple comparisons test.

### Enhanced Atg9 phosphorylation at T62 and T69 increases the autophagosome formation in glia

We next sought to determine the physiological significance of Atg9 phosphorylation at T62 and T69. Transgenic flies expressing the Atg9 variants with the threonine residues mutated to either alanine (non-phosphorylatable Atg9^T62A^, Atg9^T69A^, and Atg9^T62A-T69A^) or glutamate (phosphomimetic Atg9^T62E^, Atg9^T69E^, and Atg9^T62E-T69E^) were created. These Atg9 variants were expressed properly as examined by WB analysis (Figures S1A and S1B). Taking advantage of these variants, glial autophagosomes in the selected anterior/dorsal glia-rich region of the adult fly brain were analyzed by expressing the *UAS-mCherry.Atg8a* under the control of a pan-glial driver (*UAS-mCherry.Atg8a; repo-*GAL4, Figures 1E). Consistent with our previous findings, downregulating dAux expression increased the number of the mCherry.Atg8a-labeled autophagosomes in glia (Figures 1F and 1G). In contrast, *atg9*-RNAi expression, which efficiency was verified by qRT-PCR (Figure S1C), caused a significant reduction on the autophagosome number in glia (Figures 1H and 1I). Co-expression of *atg9*-RNAi suppressed the *daux*-RNAi-induced increase in the autophagosome number (Figures 1J and 1K). These results suggest that Atg9 acts downstream of dAux and is important for dAux-mediated autophagosome formation. Consistent to results from *daux*-RNAi, expressing Atg9 or either of the phosphomimetic Atg9 variants caused a significant increase in the glial autophagosome number, and expressing either of the three non-phosphorylatable Atg9 variants failed to cause significant difference (Figures 1L and 1N). Taken together, these results support the notion that lack of dAux increases Atg9 phosphorylation at T62 and T69, leading to enhanced number of autophagosomes in glia.

### Expression of the non-phosphorylatable Atg9 variants suppresses the increased autophagosome formation in the absence of glial dAux

To further investigate the *in-vivo* relationship between dAux and the Atg9 phosphorylation at these two residues, we co-expressed different Atg9 variants with *daux*-RNAi, to see if the change in Atg9 phosphorylation by these means affects the *daux-*RNAi-induced increase in the autophagosome number. Interestingly, expressing either of the three non-phosphorylatable Atg9 variants suppressed the increased autophagosome number in the absence of glial dAux, whereas expressing either of the three phosphomimetic Atg9 variants failed to reduce the increase significantly (Figures 1M and 1O). These results suggest that altered Atg9 phosphorylation levels at T62 and T69 regulate dAux-mediated autophagosome formation in glia.

### Atg1 presence recruits Atg9 for dAux interaction

It has been reported that Atg1 mediated Atg9 phosphorylation at multiple sites^16,26,27^. Our previous study also showed that dAux regulates Atg1 trafficking in autophagy. Next, we tested if Atg1 is involved in the dAux-mediated Atg9 regulation. Consistent to previous findings, our co-immunoprecipitation (Co-IP) analysis revealed that Atg1 interacted with Atg9, yet failed to detect Atg9 in the pull-downs of dAux when co-expressing dAux and Atg9, further suggesting that Atg9 is not a direct dAux substrate (Figures 2A and 2B). Nonetheless, Atg9 was detected in the pull-downs of dAux when co-expressing Atg1 (but not GFP), dAux, and Atg9 (Figures 2C). These results suggest that dAux, Atg1, and Atg9 form a complex and Atg1 expression is critical in recruiting Atg9 to the dAux-containing complex.

**Figure 2.**
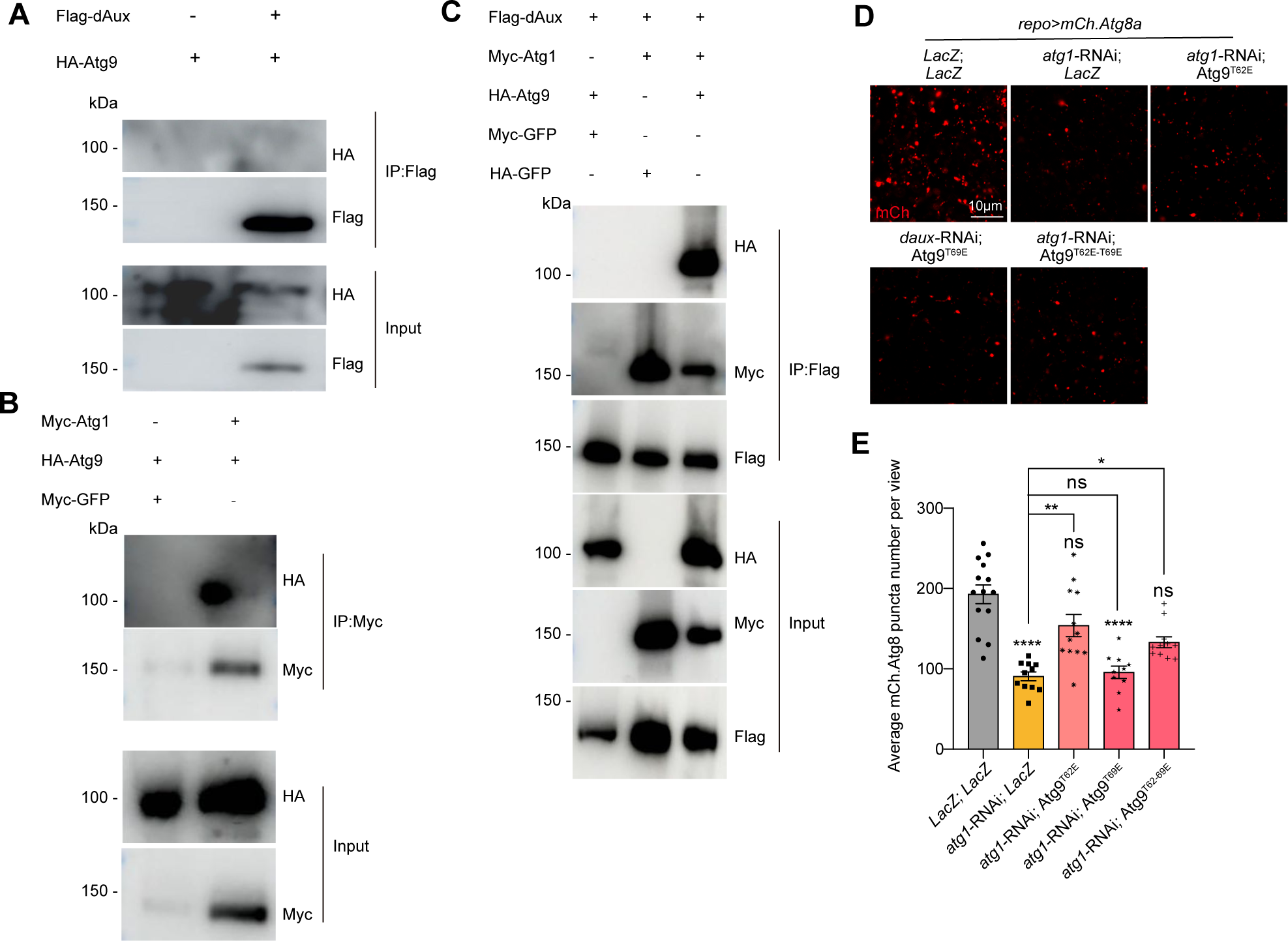
dAux-Atg9 interaction depends on Atg1. (**A**-**C**) Co-IP analysis using the fly S2 cells expressing Flag-dAux, HA-Atg9, or Myc-Atg1 in different combinations was conducted. Note that Atg9 and dAux fail to interact when co-expressing Flag-dAux and HA-Atg9 and detecting with the anti-Flag antibodies (A). Atg1 and Atg9 are within the same complex when using beads conjugated to anti-Myc antibodies to pull down Myc-Atg1 (B). Atg9 is detected in cells expressing Flag-dAux, Myc-Atg1 and HA-Atg9 pulled down by beads conjugated to anti-Flag antibodies (C). (**D** and **E**) Representative images (D) and quantifications (E) of autophagosomes in adult fly glia expressing different Atg9 variants with *atg1*-RNAi. Note that co-expression of Atg9^T62E^ and Atg9^T62E-T69E^ restores the decreased autophagosome number upon glial Atg1 depletion. Scale bars are indicated on the images. Serial confocal Z-stack sections were taken at similar planes across all genotypes, with representative images shown as a single layer. Statistical graphs are shown with scatter dots indicating the number of brain samples analyzed. Data are shown as mean ± SEM. P-values of significance (indicated with asterisks, ns no significance, * p<0.05, ** p<0.01, *** p<0.001, and **** p<0.0001) are calculated by Kruskal-Wallis tests followed by Dunn’s multiple comparisons test.

To further demonstrate a role of Atg1 in dAux-regulated Atg9 phosphorylation *in-vivo*, we co-expressed different phophomimetic Atg9 variants and analyzed their genetic relationship with Atg1. Upon *atg1*-RNAi expression, which efficiency was verified by qRT-PCR (Figure S1D), the number of autophagosomes in glia reduced. Yet, co-expression of Atg9^T62E^ or Atg9^T62E-T69E^, but not Atg9^T69E^, partially and significantly restores the decrease (Figures 2D and 2E). These results further support the notion that Atg1, which activity we have shown to be under control by dAux, functions upstream of and regulates Atg9 phosphorylation at T62 and T69.

### dAux- and Atg1-mediated Atg9 trafficking in glia depends on Atg9 phosphorylation at T62 and T69

It has been shown that Atg9 phosphorylation is involved in its trafficking to the autophagy initiation site^27–29^. We next tested whether Atg9 phosphorylation at T62 and T69, like other residues, is also important for its trafficking to autophagosomes. Consistent with the results from *daux*-RNAi, expression of Atg9 and the phosphomimetic Atg9^T62E-T69E^ increased the Atg9-Atg8a colocalization, as revealed by the dual expression of *UAS-mCherry.Atg8a* and *UAS-EGFP.Atg9* reporters in glia (Figures S2A, S2A’, and S2B). A similar increasing trend was observed for Atg9^T62E^ and Atg9^T69E^. On the contrary, the expression of non-phosphorylatable Atg9^T62A^ reduced Atg9-Atg8a colocalization significantly, and a similar decreasing trend was observed for Atg9^T69A^ and Atg9^T62A-T69A^ (Figures S2A, S2A’, and S2B). These results indicate that Atg9 trafficking to the autophagosomes is affected by Atg9 phosphorylation at T62 and T69. Notably, co-expressing either of the three non-phosphorylatable Atg9 mutants suppressed the *daux*-RNAi-induced increase in the Atg9-Atg8a colocalization (Figures 3A, 3A’, and 3B), demonstrating the importance of Atg9 phosphorylation at T62 and T69 in dAux-mediated Atg9 trafficking. Based on the requirement of Atg1 in dAux-Atg9 interaction, and also the dAux regulation on Atg1 trafficking, we also tested if Atg1 is involved in dAux-mediated Atg9 trafficking to the autophagosomes. Whereas *atg1*-RNAi expression in glia blocked Atg9 trafficking to autophagosomes, co-expression of the phosphomimetic Atg9^T62E^ or Atg9^T62E-T69E^ significantly restores the suppression, with the double variants exhibited a most potent effect (Figures 3C, 3C’, and 3D). Taken together, our results suggest that Atg9 phosphorylation at T62 and T69 contributes to the dAux-Atg1-Atg9 axis in regulating the Atg9 trafficking to the autophagosomes.

**Figure 3.**
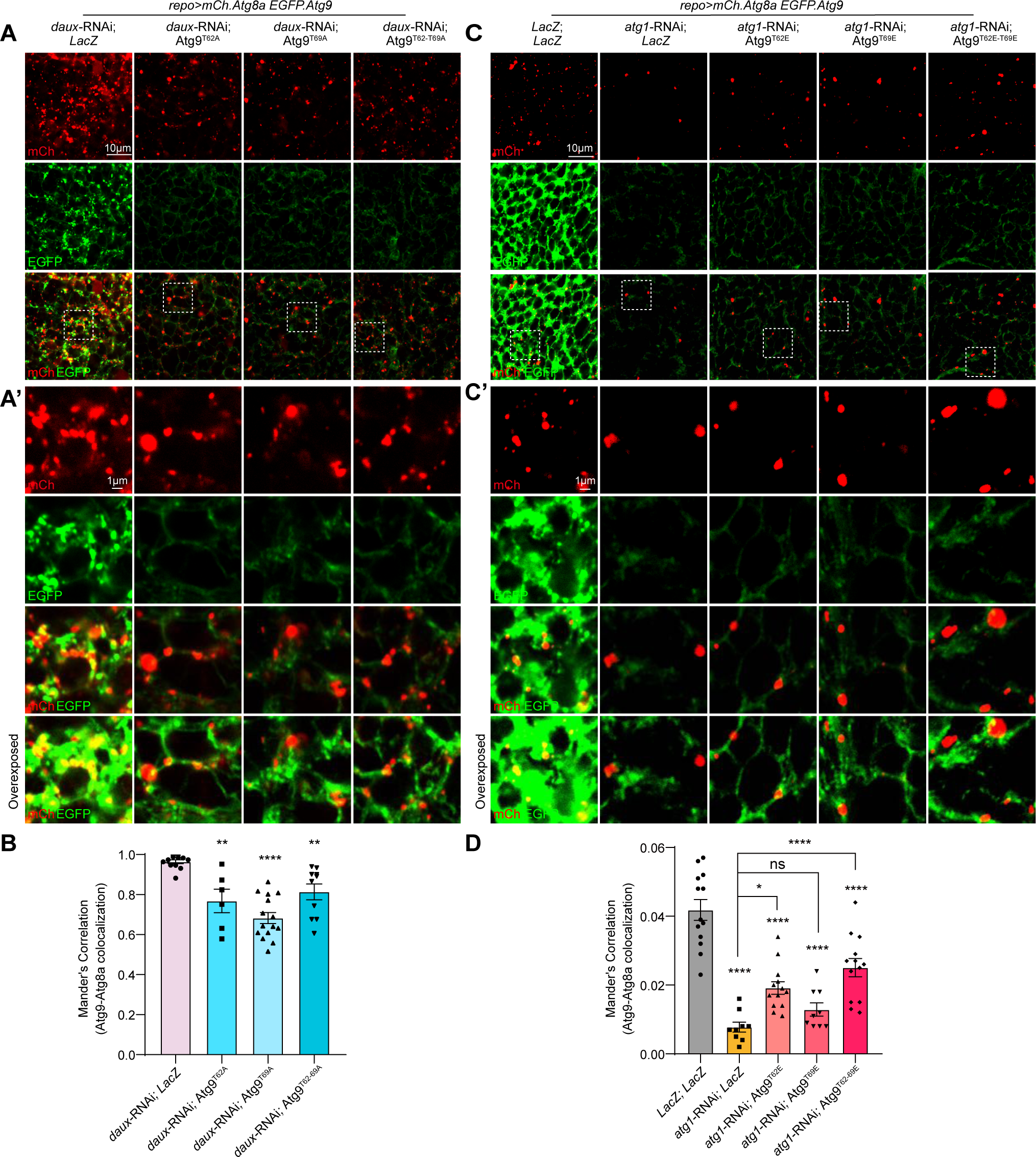
dAux- and Atg1-regulated Atg9 trafficking in glia depends on Atg9 phosphorylation at T62 and T69. (**A**-**D**) Representative images (A, A’, C, and C’) and quantifications (B and D) of Atg9-Atg8a colocalization in adult fly glia expressing different Atg9 variants with *daux*-RNAi or *atg1*-RNAi. *UAS-EGFP.Atg9* and *UAS-mCh.Atg8a* reporters were expressed in glia for analyzing the Atg9 (green)-Atg8a (red) colocalization (*UAS-mCh.Atg8a; repo-*GAL4*, UAS-EGFP.Atg9*). Note that co-expression of either of the three non-phosphorylatable Atg9 mutants partially suppresses the increase of Atg9-Atg8a colocalization upon glial dAux depletion (A, A’, and B). The Atg9-Atg8a colocalization decreases when expressing *atg1-*RNAi, which is restored by expressing Atg9^T62E^ or Atg9^T62E-T69E^ (C, C’, and D). Areas enclosed by the white dashed squares in the representative images (A and C) are enlarged in A’ and C’. An overexposed panel is listed at the bottom of A’ and C’ for better visualization of signal colocalization. Scale bars of different sizes are indicated on the images. Serial confocal Z-stack sections were taken at similar planes across all genotypes, with representative images shown as a single layer. Colocalization is analyzed using the Manders’ Correlation, taking into account the change in the protein level. Statistical graphs are shown with scatter dots indicating the number of brain samples analyzed. Data are shown as mean ± SEM. P-values of significance (indicated with asterisks, ns no significance, * p<0.05, ** p<0.01, *** p<0.001, and **** p<0.0001) are calculated by ordinary one-way ANOVA followed by Tukey’s multiple comparisons test.

### Enhanced Atg9 phosphorylation at T62 and T69 contributes to DA neurodegeneration and locomotor dysfunction

As Atg9 dysregulation has been implicated in PD^21–23^, we tested whether Atg9 phosphorylation at T62 and T69 contributes to locomotor function and DA neurodegeneration in flies. Interestingly, expression of Atg9 or either of the three phosphomimetic Atg9 variants resulted in a significant reduction in the climbing distance of 10-day-old adult male and female flies, while the expression of the non-phosphorylatable Atg9 variants failed to cause significant difference (Figure 4A). Intriguingly, co-expression of *daux*-RNAi with either of the three non-phosphorylatable Atg9 variants restored the *daux*-RNAi-induced locomotor deficits in 10-day-old adult male and female flies (Figure 4B). These results demonstrate an important role for the dAux-mediated Atg9 phosphorylation in the context of locomotor deficits implicated in PD.

**Figure 4.**
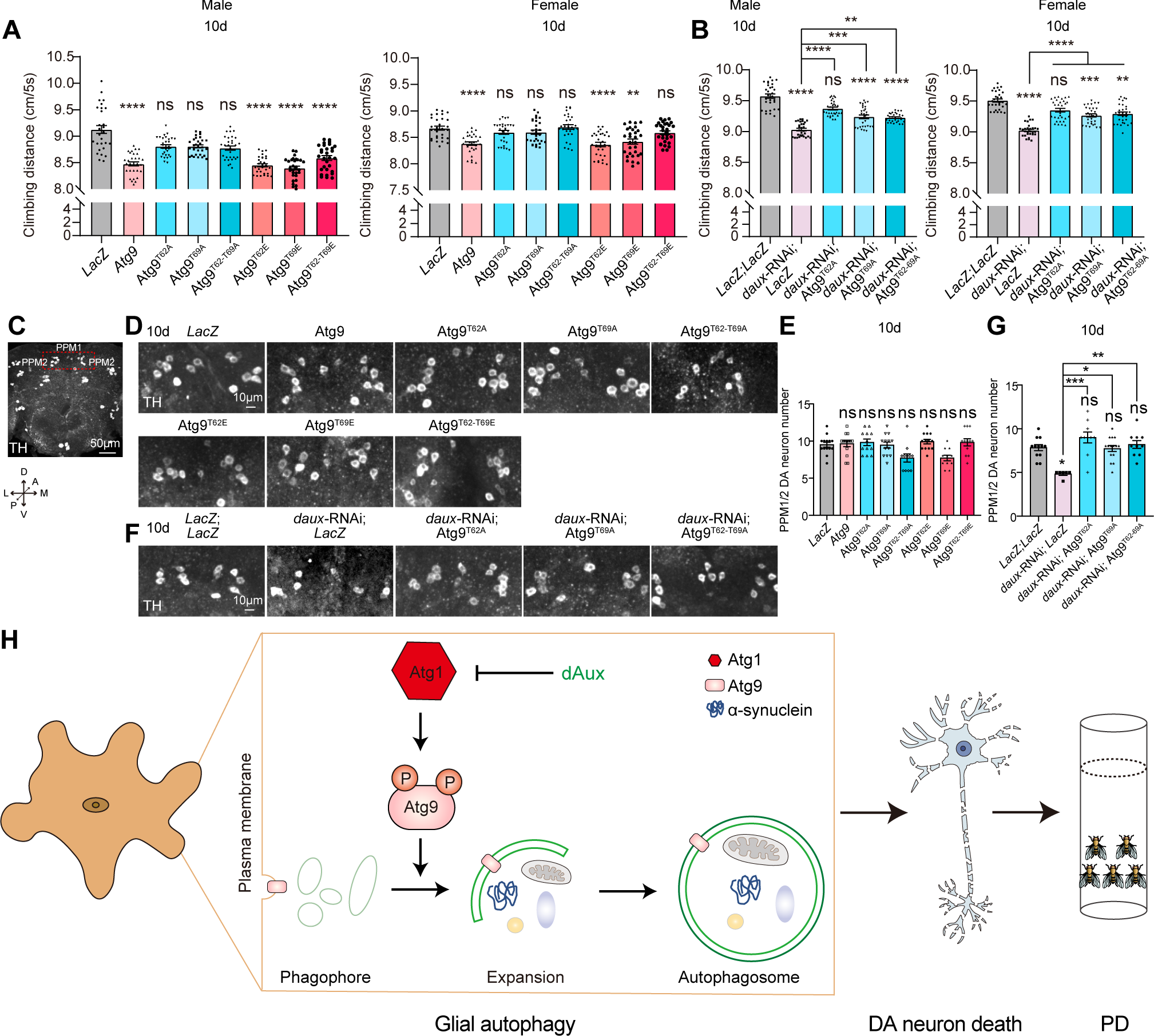
Enhanced Atg9 phosphorylation at T62 and T69 contributes to DA neurodegeneration and locomotor dysfunction. (**A** and **B**) Locomotor activity of 10-day-old adult male and female flies is analyzed. Note that expression of Atg9 or either of the three phosphomimetic Atg9 variants results in a significant reduction in the climbing distance of adult male and female flies, while the expression of the non-phosphorylatable Atg9 variants fails to cause significant difference (A). Co-expression of *daux*-RNAi with either of the three non-phosphorylatable Atg9 mutants restores the *daux*-RNAi-induced locomotor deficits in both male and female flies (B). (**C**) An acquired microscopic image of an adult fly brain stained with anti-TH antibodies revealing all DA neuron clusters, the protocerebral posterior medial (PPM)1 and PPM2 clusters enclosed by red dotted squares are selected for further analyzing. The adult fly brain is positioned with the coordinates described (A: Anterior, P: Posterior, M: Medial, L: Lateral, D: Dorsal, V: Ventral). (**D**-**G**) Representative images (D and F) and quantifications (E and G) of DA neuron number in 10-day-old adult fly brains with the indicated genotypes. Note that expression of either of the three non-phosphorylatable or phophomimetic Atg9 variants fails to cause significant changes in the DA neuron number at the PPM1/2 cluster. Co-expressing either of the three non-phosphorylatable Atg9 mutants rescues the DA neuron loss upon glial dAux depletion. (**H**) The schematic diagram illustrating a dAux-Atg1-Atg9 axis regulating glial autophagy in PD. Scale bars of different sizes are indicated on the images. Serial confocal Z-stack sections were taken at similar planes across all genotypes, with representative images shown as maximal projection. Statistical graphs are shown with scatter dots indicating the number of brain samples analyzed. In locomotion experiments, n=100 for each genotype. Data are shown as mean ± SEM. P-values of significance (indicated with asterisks, ns no significance, * p<0.05, ** p<0.01, *** p<0.001, and **** p<0.0001) are calculated by ordinary one-way ANOVA followed by Tukey’s multiple comparisons test or Kruskal-Wallis tests followed by Dunn’s multiple comparisons test.

Next, DA neurodegeneration in the 10-day-old adult fly brains was analyzed. Expressing either of the three non-phosphorylatable or phophomimetic Atg9 variants failed to cause significant changes in the DA neuron number at the PPM1/2 cluster, where DA neuron loss has been observed upon α-synuclein overexpression in flies^30^ (Figures 4C-4E). Nonetheless, *daux*-RNAi-induced DA neurodegeneration was significantly suppressed upon co-expression of either of the three non-phosphorylatable Atg9 variants (Figures 4F and 4G). These results suggest that lack of dAux-enhanced Atg9 phosphorylation at T62 and T69 contributes to DA neurodegeneration implicated in PD.

In the present study, we reported that the phosphorylation of two newly identified Atg9 residues contributes to glial autophagy implicated in PD (Figures 4H). The dAux-regulated phosphorylation at Atg9 T62 and T69 is physiologically and pathologically relevant as expressing the non-phosphorylatable or phophomimetic Atg9 variants of these two residues regulate the autophagosome formation and Atg9 trafficking to autophagosomes in adult fly glia. Our findings also suggest that Atg1 is both crucial for recruiting Atg9 to dAux-containing complex and genetically upstream in transducing the Atg9 phosphorylation signaling forward. Importantly, the dAux-Atg1-Atg9 axis of phosphorylation-dependent signaling contributes to DA neurodegeneration and locomotor dysfunction implicated in PD. Our findings further elaborate the mechanism of the autophagic degradation pathway in glia and provide insights on potential therapeutical solution for PD.

## Acknowledgements

We thank Bloomington *Drosophila* Stock Center, Vienna *Drosophila* RNAi Center, the Core Facility of *Drosophila* Resource and Technology, Shanghai Institute of Biochemistry and Cell Biology, Chinese Academy of Sciences for fly stocks, We also thank the Molecular Imaging Core Facility (MICF), the Molecular and Cell Biology Core Facility (MCBCF), and the Multi-Omics Core Facility (MOCF) at the School of Life Science and Technology, ShanghaiTech University for providing technical support; DroBot Biotechnology for quality fly food supply, delicate fly-keeping service, and experimental device design; Ho lab members for discussion and comments. This work was financially supported by grants from ShanghaiTech and National Yang Ming Chiao Tung University (intramural), National Natural Science Foundation of China (32170962), 2030 Cross-Generation International Outstanding Young Scholars Program National Science and Technology Council Taiwan (113-2628-B-A49-007), and Brain Research Center National Yang Ming Chiao Tung University from The Featured Areas Research Center Program within the framework of the Higher Education Sprout Project by the Ministry of Education (MOE) in Taiwan.

## Author contributions

L.W, S.Y, S.Z, and M.S.H conceived and designed the study. L.W, S.Y, S.Z, and H.W performed the experiments. L.W, S.Y, S.Z, Y.T, and M.S.H analyzed the data and wrote the paper. All authors read and approved the manuscript.

