## Supplementary Materials for "New Atg9 phosphorylation sites regulate autophagic trafficking in glia"

**This PDF file includes:**

Materials and Methods

Supplementary Figures S1-S2

Table S1

#### Materials and Methods

##### *Drosophila* genetics

All fly crosses were conducted at 25°C. Strains are acquired from Bloomington *Drosophila* Stock Center, Vienna *Drosophila* RNAi Center, or used previously: *UAS-LacZ* (BL#1777), *UAS-mCherry.Atg8a* (BL#37750), *UAS-atg9-RNAi* (BL#28055), *UAS-atg1*-RNAi (BL#80434); *UAS-daux*-RNAi (V#16182), *repo-GAL4*[1], and *UAS-EGFP.Atg9*[2]. Adult flies at 10-day-old were analyzed. Detailed genotypes are listed in Table S1.

##### Plasmid cloning

The DNA sequences expressing Atg9 and Atg9 mutants (Atg9$^{T62A}$, Atg9$^{T69A}$, Atg9$^{T62A-T69A}$, Atg9$^{T62E}$, Atg9$^{T69E}$, and Atg9$^{T62E-T69E}$) were synthetized and then subcloned into *pUAST-attB* vectors containing 3xHA epitope tag by Genscript (Nanjing, China). Fly embryo microinjection was carried out by the *Drosophila* Core Facility, Institute of Biochemistry and Cell Biology, Chinese Academy of Sciences.

##### qRT-PCR

Total RNAs were extracted using adult fly heads by TransZol Up (Cat. #ET111-01, TransGen, Beijing, China), and then revise transcription with HiScript III RT SuperMix (Cat. #R323-01, Vazyme, Nanjing, China). qRT-PCR reactions were performed using ChamQ Universal SYBR qPCR Master Mix (Cat. #Q711-02, Vazyme) and ABI 7500 RT-PCR system. The comparative Threshold Cycle (Ct) method was used for quantification. The Ct values were normalized to *rp49*. Relative quantification was performed using the ΔΔCT method.

Primers used are listed below:

*atg1* is described previously[3].

*rp49*-F: CCACCAGTCGGATCGATATGC

*rp49*-R: CTCTTGAGAACGCAGGCGACC

*atg9*-F: AGCAGAAGCACGGATTCACA

*atg9*-R: GCAGTGCATCACAAAGGCAA

**Immunohistochemistry**

Adult fly brains were dissected and fixed using 4% formaldehyde for 40 minutes, and then washed 3 times with PBT (PBS + 0.1% TX-100). The brains were blocked in PBT with 5% normal donkey serum, and stained with primary antibodies at 4°C overnight, and then secondary antibodies at room temperature for 2 hours. Primary antibody used is rat anti-TH (1:300, NB300-109, Novus Biologicals, Littleton, CO, USA). The secondary antibody used were from Jackson ImmunoResearch (West Grove, PA, USA) is donkey anti-rat Cy3 (1:1000, Cat. #712-165-153).

**Western blot analysis**

A motorised pestle (Cat. #116005500, MP Biomedicals, Irvine, CA, USA) was used to homogenise adult fly heads in lysis buffer (0.4% NP-40, 20% glycerol, 0.2 mM EDTA, 100 mM Tris-HCl pH7.5, 150 mM NaCl, 0.5 mM phosphodiesterase inhibitors, 2%

Tween 20, and 1 mM PMSF). Proteins were separated on SDS-PAGE gels and transferred to PVDF membranes (Cat. #IPFL00010, Millipore, Billerica, MA, USA). The membranes were blocked in 5% fat-free milk with PBST for 40 minutes. Samples were incubated with the primary antibodies at 4°C overnight, and then HRP-conjugated secondary antibodies at room temperature for 2 hours. Primary antibodies used include: mouse anti-α-Tubulin (1:5000, Cat. #T9026, Sigma), rabbit anti-HA (1:1000, Cat. #3724T, Cell Signaling Technology, CST, Danvers, MA, USA), rabbit-anti-Myc (1:2000, Cat. #0912-2, Hua An Biotechnology), and mouse anti-Flag (1:1000, Cat. #F3165, Sigma, St. Louis, MO, USA). Secondary antibodies used are from Jackson ImmunoResearch: goat anti-mouse-HRP (1:5000, Cat. #115-035-003) and goat anti-rabbit-HRP (1:5000, Cat. #111-036-003). Clarity Western ECL Substrate (Cat. #SB-WB001, share-bio) was used to visualize bands.

**Co-immunoprecipitation**

*Drosophila* S2 cells were cultured at 28°C with Schneider's medium (Cat. #21720024, Gibco), and the Effectene Transfection Reagent (Cat. #301425, Qiagen) was used for transfection. Cells were lysed on ice using lysis buffer for 30 minutes. For pull-downs, the samples were centrifuged at 13,000 rpm at 4°C for 10 minutes. Prewashed anti-FLAG(R) M2 beads (Cat. #A2220; Sigma) or rabbit-anti-Myc (1:2000, Cat. #0912-2, Hua An Biotechnology) were used to incubate the soluble supernatant at 4°C overnight. After that, samples were washed with lysis buffer for 30 minutes three times and then used for western blot analysis.

**Fly locomotion**

The rapid iterative negative geotaxis (RING) assay was used as previously described[4-6]. Flies were collected and placed in vials with fly food (no yeast) for 1 day before transferring into the cylinders for testing. 100 flies per genotype were analyzed in ten cylinders (inner diameter: 20 mm; height: 140 mm), and each contains ten unisex flies. An initial mechanical shock was applied six times to tap down all flies to the bottom of the cylinder. Climbing distances were measured and averaged by a self-designed RflyDetection software, allowing automatic detection of the fly position within the cylinder using video images recorded every 5 seconds (Sony digital camera, HDR-CX220E). At least three independent experiments were performed.

**Confocal microscopy and quantification**

Images were acquired by scanning a serial Z-stack sections at the similar plane using Nikon C2 or TI2-E+CSU W1 Sora Spinning Disk confocal microscope (20x objective NA=0.75, 60x oil objective NA=1.4, Tokyo, Japan). Representative single layer images or maximum projection images are shown. ImageJ (National Institutes of Health) was used to analyze puncta. The intensity threshold was used to quantify numbers of dots. Colocalization plugin was used for colocalization measurements, and the results were shown as Manders' Correlation (M1 or M2).

**Statistical analysis**

GraphPad Prism 8 (Dotmatics, Boston, Massachusetts, USA) was used to present data and analyze significance. Shapiro-Wilk normality test was firstly used to check the data normal distribution, two-tailed unpaired t-test or ordinary one-way ANOVA followed by Tukey's multiple comparisons test was used for normally distributed datasets, Mann-Whitney test or Kruskal-Wallis tests followed by Dunn's multiple comparisons test were used for datasets without a normal distribution. P value less than 0.05 is considered significant. ns: no significance, $p \geq 0.05$; *: $p<0.05$; **: $p<0.01$; ***: $p<0.001$; ****: $p<0.0001$.

Figure S1

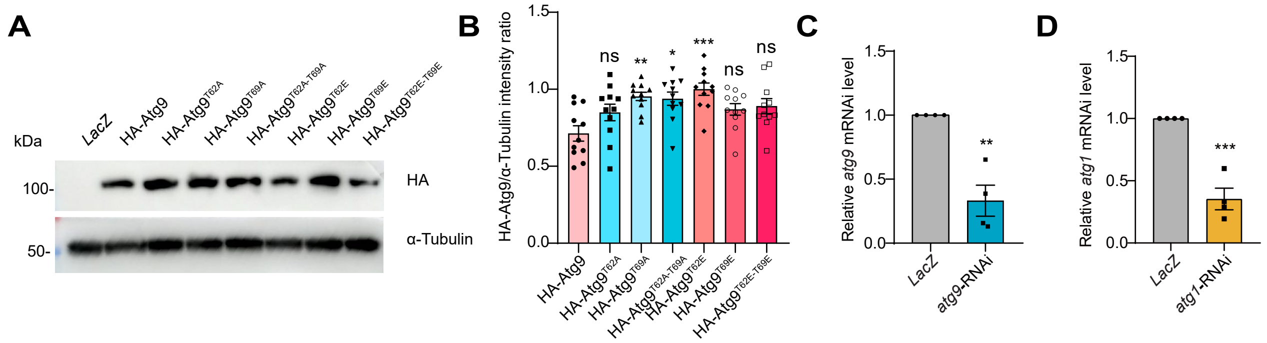
A
B
C
D
kDa
LacZ
HA-Atg9
HA-Atg9T62A
HA-Atg9T69A
HA-Atg9T62A-T69A
HA-Atg9T62E
HA-Atg9T69E
HA-Atg9T62E-T69E
100-
50-
HA
α-Tubulin
HA-Atg9/α-Tubulin intensity ratio
ns
**
*
***
ns
ns
Relative atg9 mRNAi level
LacZ
atg9-RNAi
**
Relative atg1 mRNAi level
LacZ
atg1-RNAi
***

**Figure S1 (related to Figures 1 and 2). Verification of different transgenic fly lines used in the study**

(**A** and **B**) Atg9 variants with the threonine residues mutated to either alanine ($Atg9^{T62A}$, $Atg9^{T69A}$, and $Atg9^{T62A\text{-}T69A}$) or glutamate ($Atg9^{T62E}$, $Atg9^{T69E}$, and $Atg9^{T62E\text{-}T69E}$) were constructed and expressed properly in flies. (**C** and **D**) RNAi targeting *atg9* (C) or *atg1* (D) efficiently silenced the gene expression as verified by the qRT-PCR analysis. Statistical graphs are shown with scatter dots indicating the number of biological replicates analyzed. Data are shown as mean ± SEM. P-values of significance (ns: no significance, $p \geq 0.05$; *: $p<0.05$; **: $p<0.01$; ***: $p<0.001$; ****: $p<0.0001$) are calculated by two-tailed unpaired t-test or ordinary one-way ANOVA followed by Tukey's multiple comparisons test.

### Figure S2

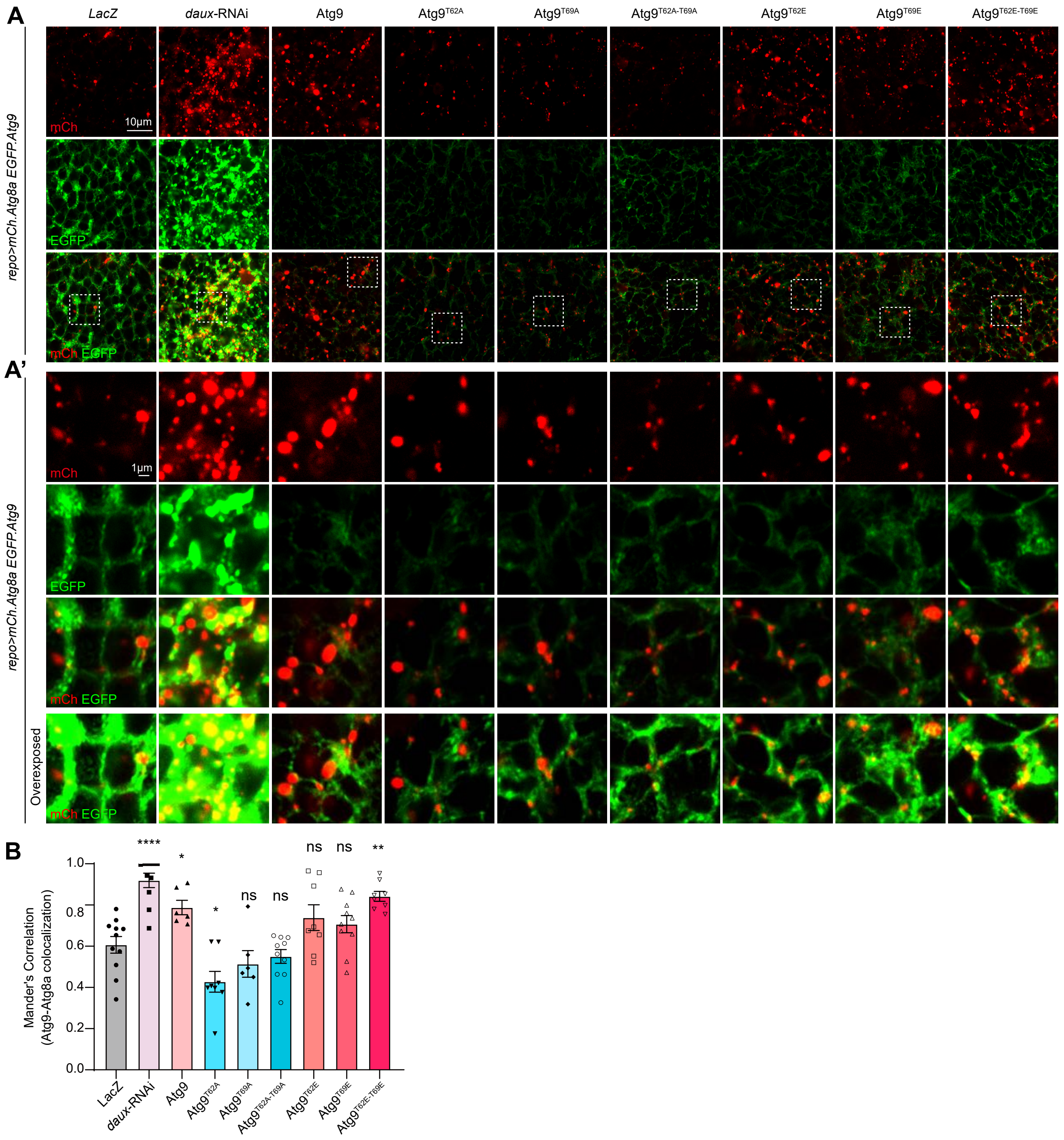
A
LacZ
daux-RNAi
Atg9
Atg9T62A
Atg9T69A
Atg9T62A-T69A
Atg9T62E
Atg9T69E
Atg9T62E-T69E
repo>mCh.Atg8a EGFP.Atg9
mCh
10µm
EGFP
mCh EGFP
A'
1µm
Overexposed
B
Mander's Correlation
(Atg9-Atg8a colocalization)
1.0
0.8
0.6
0.4
0.2
0.0
****
*
*
ns
ns
ns
ns
**
LacZ
daux-RNAi
Atg9
Atg9T62A
Atg9T69A
Atg9T62A-T69A
Atg9T62E
Atg9T69E
Atg9T62E-T69E

**Figure S2 (related to Figure 3). The altered phosphorylation at T62 and T69 regulates Atg9 trafficking to the autophagosomes**

(**A** and **B**) Representative images (A and A') and quantifications (B) of Atg9-Atg8a colocalization in adult fly glia expressing *daux*-RNAi or different Atg9 variants. *UAS-EGFP.Atg9* and *UAS-mCh.Atg8a* reporters were expressed in glia for analyzing the Atg9 (green)-Atg8a (red) colocalization (*UAS-mCh.Atg8a; repo*-GAL4, *UAS-EGFP.Atg9*). Note that the Atg9-Atg8a colocalization increases when expressing *daux*-RNAi, Atg9, or phophomimetic Atg9$^{T62E\text{-}T69E}$, and decreases when expressing non-phosphorylatable Atg9$^{T62A}$. Areas enclosed by the white dashed squares in the representative images (A) are enlarged in A'. An overexposed panel is listed at the bottom of A' for better visualization of signal colocalization. Scale bars of different sizes are indicated on the images. Serial confocal Z-stack sections were taken at similar planes across all genotypes, with representative images shown as a single layer. Colocalization is analyzed using the Manders' Correlation, taking into account the change in the protein level. Statistical graphs are shown with scatter dots indicating the number of brain samples analyzed. Data are shown as mean ± SEM. P-values of significance (indicated with asterisks, ns no significance, * $p<0.05$, ** $p<0.01$, *** $p<0.001$, and **** $p<0.0001$) are calculated by ordinary one-way ANOVA followed by Tukey's multiple comparisons test.

**Table S1**

Detailed fly genotypes in each experiment categorized by Figures.

| Figure | Genotype |
|---|---|
| Fig. 1 | |
| (A-D) | *repo*-GAL4/*UAS-LacZ* |
| | *UAS-daux*-RNAi/+; *repo*-GAL4/+ |
| (E) | *UAS-mCherry.Atg8a/+; repo-GAL4/+* |
| (F and G) | *UAS-mCherry.Atg8a/+; repo-GAL4/UAS-LacZ* |
| | *UAS-mCherry.Atg8a/UAS-daux*-RNAi*; repo-GAL4/+* |
| (H and I) | *UAS-mCherry.Atg8a/+; repo-GAL4/UAS-LacZ* |
| | *UAS-mCherry.Atg8a/+; repo-GAL4/UAS-atg9*-RNAi |
| (J and K) | *UAS-mCherry.Atg8a/UAS-LacZ; repo-GAL4/UAS-LacZ* |
| | *UAS-mCherry.Atg8a/UAS-daux*-RNAi*; repo-GAL4/UAS-LacZ* |
| | *UAS-mCherry.Atg8a/UAS-daux*-RNAi*; repo-GAL4/UAS-atg9*-RNAi |
| (L and N) | *UAS-mCherry.Atg8a/+; repo-GAL4/UAS-LacZ* |
| | *UAS-mCherry.Atg8a/+; repo-GAL4/UAS-HA-Atg9* |
| | *UAS-mCherry.Atg8a/+; repo-GAL4/UAS-HA-*$Atg9^{T62A}$ |
| | *UAS-mCherry.Atg8a/+; repo-GAL4/UAS-HA-*$Atg9^{T69A}$ |
| | *UAS-mCherry.Atg8a/+; repo-GAL4/UAS-HA-*$Atg9^{T62A\text{-}T69A}$ |
| | *UAS-mCherry.Atg8a/+; repo-GAL4/UAS-HA-*$Atg9^{T62E}$ |
| | *UAS-mCherry.Atg8a/+; repo-GAL4/UAS-HA-*$Atg9^{T69E}$ |
| | *UAS-mCherry.Atg8a/+; repo-GAL4/UAS-HA-*$Atg9^{T62E\text{-}T69E}$ |
| (M and O) | *UAS-mCherry.Atg8a/UAS-LacZ; repo-GAL4/UAS-LacZ* |
| | *UAS-mCherry.Atg8a/UAS-daux*-RNAi*; repo-GAL4/UAS-LacZ* |
| | *UAS-mCherry.Atg8a/UAS-daux*-RNAi*; repo-GAL4/UAS-HA-*$Atg9^{T62A}$ |
| | *UAS-mCherry.Atg8a/UAS-daux*-RNAi*; repo-GAL4/UAS-HA-*$Atg9^{T69A}$ |
| | *UAS-mCherry.Atg8a/UAS-daux*-RNAi*; repo-GAL4/UAS-HA-*$Atg9^{T62A\text{-}T69A}$ |
| | *UAS-mCherry.Atg8a/UAS-daux*-RNAi*; repo-GAL4/UAS-HA-*$Atg9^{T62E}$ |
| | *UAS-mCherry.Atg8a/UAS-daux*-RNAi*; repo-GAL4/UAS-HA-*$Atg9^{T69E}$ |
| | *UAS-mCherry.Atg8a/UAS-daux*-RNAi*; repo-GAL4/UAS-HA-*$Atg9^{T62E\text{-}T69E}$ |
| Fig. 2 | |
| (D and E) | *UAS-mCherry.Atg8a/UAS-LacZ; repo-GAL4/UAS-LacZ* |
| | *UAS-mCherry.Atg8a/UAS-atg1*-RNAi*; repo-GAL4/UAS-LacZ* |
| | *UAS-mCherry.Atg8a/UAS-atg1*-RNAi*; repo-GAL4/UAS-HA-*$Atg9^{T62E}$ |
| | *UAS-mCherry.Atg8a/UAS-atg1*-RNAi*; repo-GAL4/UAS-HA-*$Atg9^{T69E}$ |
| | *UAS-mCherry.Atg8a/UAS-atg1*-RNAi*; repo-GAL4/UAS-HA-*$Atg9^{T62E\text{-}T69E}$ |
| Fig. 3 | |
| (A and B) | *UAS-mCherry.Atg8a/UAS-daux*-RNAi*; UAS-EGFP-Atg9, repo-GAL4/UAS-LacZ* |

| | |
|---|---|
| | *UAS-mCherry.Atg8a/UAS-daux*-RNAi*; UAS-EGFP-Atg9, repo-GAL4/UAS-HA-Atg9$^{T62A}$* |
| | *UAS-mCherry.Atg8a/UAS-daux*-RNAi*; UAS-EGFP-Atg9, repo-GAL4/UAS-HA-Atg9$^{T69A}$* |
| | *UAS-mCherry.Atg8a/UAS-daux*-RNAi*; UAS-EGFP-Atg9 repo-GAL4/UAS-HA-Atg9$^{T62A-T69A}$* |
| (C and D) | *UAS-mCherry.Atg8a/UAS-LacZ; UAS-EGFP-Atg9, repo-GAL4/UAS-LacZ* |
| | *UAS-mCherry.Atg8a/UAS-atg1*-RNAi*; UAS-EGFP-Atg9, repo-GAL4/UAS-LacZ* |
| | *UAS-mCherry.Atg8a/UAS-atg1*-RNAi*; UAS-EGFP-Atg9, repo-GAL4/UAS-HA-Atg9$^{T62E}$* |
| | *UAS-mCherry.Atg8a/UAS-atg1*-RNAi*; UAS-EGFP-Atg9, repo-GAL4/UAS-HA-Atg9$^{T69E}$* |
| | *UAS-mCherry.Atg8a/UAS-atg1*-RNAi*; UAS-EGFP-Atg9 repo-GAL4/UAS-HA-Atg9$^{T62E-T69E}$* |
| Fig. 4 | |
| (A, D, and E) | *repo-GAL4/UAS-LacZ* |
| | *repo-GAL4/UAS-HA-Atg9* |
| | *repo-GAL4/UAS-HA-Atg9$^{T62A}$* |
| | *repo-GAL4/UAS-HA-Atg9$^{T69A}$* |
| | *repo-GAL4/UAS-HA-Atg9$^{T62A-T69A}$* |
| | *repo-GAL4/UAS-HA-Atg9$^{T62E}$* |
| | *repo-GAL4/UAS-HA-Atg9$^{T69E}$* |
| | *repo-GAL4/UAS-HA-Atg9$^{T62E-T69E}$* |
| (B, F, and G) | *UAS-LacZ/+; repo-GAL4/UAS-LacZ* |
| | *UAS-daux*-RNAi/+*; repo-GAL4/UAS-LacZ* |
| | *UAS-daux*-RNAi/+*; repo-GAL4/UAS-HA-Atg9$^{T62A}$* |
| | *UAS-daux*-RNAi/+*; repo-GAL4/UAS-HA-Atg9$^{T69A}$* |
| | *UAS-daux*-RNAi/+*; repo-GAL4/UAS-HA-Atg9$^{T62A-T69A}$* |
| Fig. S1 | |
| (A and B) | *repo-GAL4/UAS-LacZ* |
| | *repo-GAL4/UAS-HA-Atg9* |
| | *repo-GAL4/UAS-HA-Atg9$^{T62A}$* |
| | *repo-GAL4/UAS-HA-Atg9$^{T69A}$* |
| | *repo-GAL4/UAS-HA-Atg9$^{T62A-T69A}$* |
| | *repo-GAL4/UAS-HA-Atg9$^{T62E}$* |
| | *repo-GAL4/UAS-HA-Atg9$^{T69E}$* |
| | *repo-GAL4/UAS-HA-Atg9$^{T62E-T69E}$* |
| (C) | *repo-GAL4/UAS-LacZ* |
| | *repo-GAL4/UAS-atg9*-RNAi |

| | |
|---|---|
| (D) | *repo-GAL4/UAS-LacZ* |
| | *UAS-atg1*-RNAi/+; *repo-GAL4/+* |
| Fig. S2 | |
| (A and B) | *UAS-mCherry.Atg8a/+; UAS-EGFP-Atg9, repo-GAL4/UAS-LacZ* |
| | *UAS-mCherry.Atg8a/UAS-daux*-RNAi*; UAS-EGFP-Atg9, repo-GAL4/+* |
| | *UAS-mCherry.Atg8a/+; UAS-EGFP-Atg9, repo-GAL4/UAS-HA-Atg9* |
| | *UAS-mCherry.Atg8a/+; UAS-EGFP-Atg9, repo-GAL4/UAS-HA-* $Atg9^{T62A}$ |
| | *UAS-mCherry.Atg8a/+; UAS-EGFP-Atg9, repo-GAL4/UAS-HA-* $Atg9^{T69A}$ |
| | *UAS-mCherry.Atg8a/+; UAS-EGFP-Atg9, repo-GAL4/UAS-HA-* $Atg9^{T62A\text{-}T69A}$ |
| | *UAS-mCherry.Atg8a/+; UAS-EGFP-Atg9, repo-GAL4/UAS-HA-* $Atg9^{T62E}$ |
| | *UAS-mCherry.Atg8a/+; UAS-EGFP-Atg9, repo-GAL4/UAS-HA-* $Atg9^{T69E}$ |
| | *UAS-mCherry.Atg8a/+; UAS-EGFP-Atg9, repo-GAL4/UAS-HA-* $Atg9^{T62E\text{-}T69E}$ |
